# A convenient and simple system for *in vivo* ^13^CO_2_-labelling in plant leaves: Enhancing accessibility for tracer studies in photosynthesis

**DOI:** 10.1101/2025.03.18.643862

**Authors:** Anish Kaachra, Mamta, Shiv Shanker Pandey

## Abstract

Tracer experiments using ^13^CO_2_ is a valuable tool for studying the metabolic flux of carbon during photosynthesis, providing precise insights into carbon assimilation and distribution within plant systems. However, the complexity of designing and setting up a ^13^CO_2_ delivery system is a significant factor limiting the use of tracer experiments in many research laboratories. In the present work, we report a simple ^**13**^CO_2_ delivery system that can be used in conjunction with an Infrared gas analyser (IRGA) for short duration in vivo ^13^C labelling in plant leaves. Briefly, ^**13**^CO_2_ is released in an external airtight container upon acidification of NaH^13^CO_3_ and subsequently introduced into the leaf chamber of an IRGA. A defined concentration of ^13^CO_2_ (± 60 ppm for about 2 minutes) can be generated by using an appropriate volume and concentration of NaH^13^CO_3_ and HCl. The functionality of the ^13^CO_2_ labelling system was assessed by feeding ^13^CO_2_ to the tobacco leaf for two different time intervals (30 secs and 2 min) under controlled environment conditions. The amino acids fractions from both labelled and unlabelled control plants were analysed using liquid chromatography-mass spectrometry (LC-MS) followed by determination of changes in ^13^C enrichment. The results showed that after 30 seconds, only two amino acids, serine and alanine, exhibited a significant increase in ^13^C enrichment, with values of 8.9% and 8.3%, respectively, compared to the unlabelled control. In contrast, after 2 minutes, a significant increase in ^13^C enrichment was observed in serine (24.1%), glycine (13.8%), aspartate (13.5%), and alanine (25.9%) as compared to the unlabelled control. The results, therefore, confirmed the workability of the system for feeding ^13^CO_2_ to a plant leaf in a time dependent manner. In addition to its simpler configuration and better control over environmental conditions, the major advantage of this system is its portability, allowing users to conduct ^13^CO_2_ experiments on plants growing in their natural habitat while minimizing physiological and metabolic alterations.

## Introduction

Enhancing photosynthesis or the efficiency with which plants assimilate atmospheric CO_2_ has long been a key focus for boosting crop yields (Smith et al., 2023: Li et al., 2023). Over the years, numerous efforts have been undertaken and continue to this day to improve photosynthetic efficiency in plants (Garcia et al., 2022). Additionally, plant biologists are exploring superior photosynthetic variants across diverse agro-climatic regions (Hussain et al., 2021; Ort et al., 2022). The major research efforts, such as enhancing Rubisco kinetics, incorporating the C_4_ pathway into C_3_ plants, and reducing photorespiration in C_3_ plants, are all fundamentally focused on improving plant photosynthesis (Croce et al., 2024; Suboktagin, S. et al., 2024). Notably, these studies call for robust and reliable molecular, biochemical and physiological investigations to assess any changes in photosynthetic efficiency. Among different physiological analyses, gas exchange measurements for CO_2_ and O_2_ using infrared gas analysers (IRGAs), chlorophyll fluorescence analysis, pulse-amplitude-modulation (PAM) fluorometry, and hyperspectral imaging are routinely employed in photosynthesis measurements. In recent times, isotope labelling has emerged as a powerful tool offering detailed understanding into the dynamics of carbon and oxygen within the photosynthetic process (Walker et al., 2024; Westhoff and Weber, 2024). The use of stable isotopes like carbon-13 (^13^C) and oxygen-18 (^18^O) allows researchers to track the movement of these elements through metabolic pathways, providing deeper insights into carbon fixation and oxygen evolution. Combined with advanced analytical methods like mass spectrometry and nuclear magnetic resonance (NMR), isotope labelling is valuable for studying isotopic discrimination, flux analysis, and metabolic regulation under diverse environmental conditions (Westhoff and Weber, 2024).

Tracer studies with ^13^CO_2_ feeding have been pivotal in advancing our knowledge of fundamental biological processes in plants. These studies contributed greatly to understand various aspects of plant carbon metabolism, such as the dynamics of carbon allocation and storage across tissues, the diurnal regulation of photosynthate distribution, carbon use efficiency under stress conditions, and the carbon dynamics governing source-sink relationships. Notably,^13^CO_2_ tracing has been used in several intriguing inquiries including determination of metabolic origin of non-photorespiratory CO_2_ release during photosynthesis (Xu et al., 2021), defining flux of photorespiratory intermediates (Abadie and Tcherkez, 2021), identifying impaired efficiency of C_4_ photosynthesis in maize under low irradiance (Medeiros et al., 2022), and elucidating distinct patterns of photosynthetic carbon flux between algae, C_3_ and C_4_ plants (Treves et a., 2021). In addition, ^13^C labelling finds a great application to study the impact of genetic modifications on carbon metabolism and partitioning in engineered plants. Despite this recognised importance of ^13^CO_2_ tracer experiments, this technique is not as widely used across different laboratories, especially when compared to other physiological methods for studying photosynthesis. The complexity involved in designing and setting up an efficient ^13^CO_2_ delivery system could possibly be one reason for the limited adoption of this technique. Key factors for an efficient ^13^CO_2_ delivery include precise control of gas concentration, uniform distribution, non-disruption to plant growth, rapid quenching of tissue after feeding and cost-effectiveness. Additionally, it is important to maintain consistent environmental conditions, such as temperature, light intensity, and humidity, while feeding ^13^CO_2_ to the plant.

There are studies where specially designed systems have been successfully utilised for feeding ^13^CO_2_ to the plant leaves. Examples include *in vivo* ^13^C-labelling system for determination of rate-limiting steps of the C_3_ photosynthetic pathway in *Nicotiana tabacum* leaves (Hasunuma et al., 2010), labelling systems to resolve intracellular fluxes of the C metabolism in illuminated intact *Arabidopsis thaliana* rosettes (Szecowka et al., 2013; Heise et al., 2014), a setup for studying ^13^CO_2_ labeling kinetics in maize (Medeiros et al., 2022; Arrivault et al., 2017), among others. It is important to note that all of these reported systems require ^13^CO_2_ cylinder, mass flow controller, humidifier, gas analyser, temperature controller, and a mechanism for rapid quenching the leaf after ^13^CO_2_ delivery. While all these components provide precise control over the rate and concentration of ^13^CO_2_ delivery, the complexity of setting up the system with these components often becomes a key factor limiting the use of ^13^CO_2_ tracing experiments across different laboratories. Another significant limitation of these systems is that, with all these components, ^13^CO_2_ delivery is restricted to the laboratory environment only. Taking all of this into account, we developed an alternative ^13^CO_2_ delivery system with a minimalist configuration that offers improved accessibility and portability. The Infra-Red Gas Analyzer (IRGA) is a versatile tool commonly used in plant physiology research to measure different photosynthetic parameters. In our system, an external container is connected to the IRGA inlet via tubing, allowing the ^13^CO_2_ released inside the container to flow into the leaf chamber, where the plant leaf being studied is placed. The release of ^13^CO_2_ is achieved by acidifying NaH^13^CO_3_ with HCl. By using a specific volume and concentration of NaH^13^CO_3_ and HCl, a defined concentration of ^13^CO_2_ (± 60 ppm) can be delivered to the leaf chamber for around 2 minutes. Along with its simpler configuration, using IRGA for ^13^CO_2_ delivery allows greater control over different environmental parameters, such as humidity, leaf temperature, and light intensity, while feeding ^13^CO_2_ to the plant leaf. Another key advantage is the system’s portability, allowing ^13^CO_2_ feeding to be performed on plants growing in their natural habitat without the need for transplantation which could potentially alter their physiology or metabolism. In the present study, we configured the existing IRGA instrument to feed ^13^CO_2_ to tobacco leaves for 30 secs and 2 min under controlled environment conditions. Mass spectrometry (LC-MS) based analysis of ^13^C enrichment revealed higher levels of ^13^C in amino acids after 2 min of ^13^CO_2_ feeding as compared to 30 seconds. These results demonstrate the system’s workability, especially for studies aiming to provide comparative modulation of photosynthetic CO_2_ flux.

## Material and Method

### Plant material

*Nicotiana benthamiana* plants were grown in a potting mixture (vermiculite: coco peat: perlite; 2:1:1) supplemented with half-strength of MS media (Murasshige and Skoog) at 24°C with a light/dark cycle of 16 h/ 8 h (photosynthetic photon flux density, 250 μmol m^−2^ s^−1^) and 70% relative humidity (RH).

### ^13^CO_2_ feeding system

The ^13^CO_2_ feeding system comprised of an external airtight container (diameter: 10.5 cm; length: 7.5 cm), which was connected to the inlet of the IRGA, LI-6400 (Li-COR, USA). Inside the container, a small Petri dish with a magnetic bead was placed. A 5 mL syringe was fitted into the top cover of the container, as shown in **Fig. 1**. The Petri dish holds Na_2_^13^CO3, and the syringe was used to add HCl onto the plate. The spinning of the magnetic bead on the stirrer plate ensured uniform mixing to release ^13^CO_2_ **(Fig 2)**. It is crucial to make sure that no ^13^CO_2_ escapes from the top cover or through the connecting tubing.

**Fig. 1.**
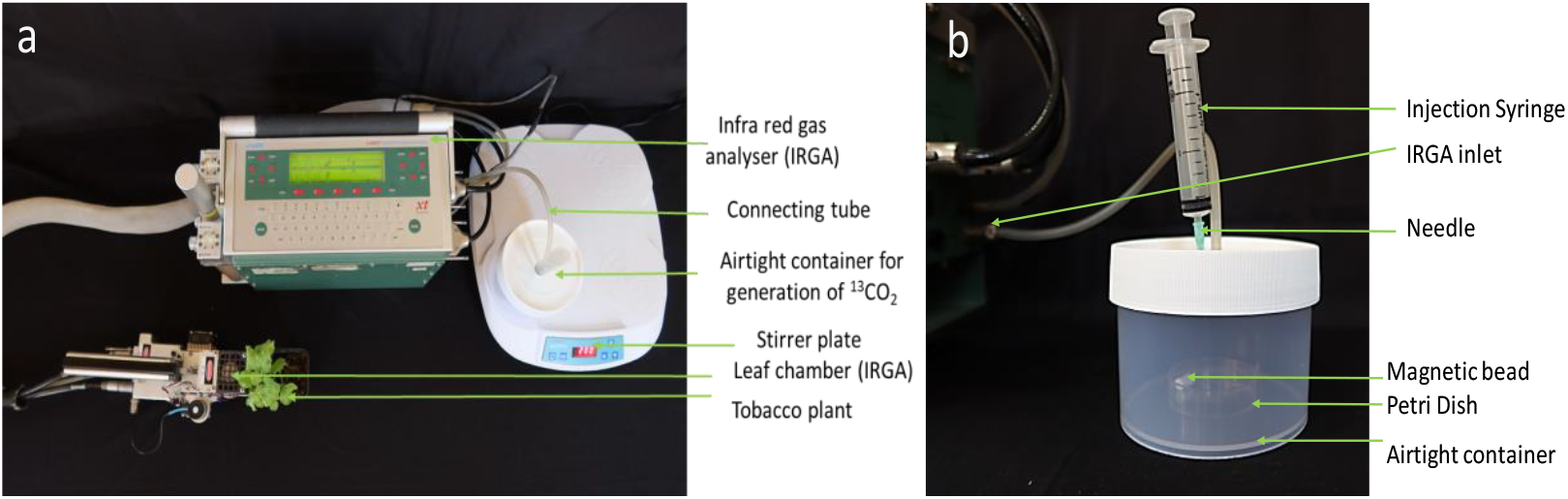
Picture showing the system used for the delivery of ^13^CO_2_ to the tobacco leaf. **(a)** Inlet of Infra-red gas analyser is connected to the airtight container used for the generation of ^13^CO_2_ through connecting tubing. The magnetic stirrer plate facilitates rotation of magnetic bead inside the container **(b)** The air tight container harbouring a petri dish with magnetic bead for uniform mixing of NaH^13^CO_3_ with HCl as detailed in the text. The injection needle is used to add HCl onto the Petri dish containing Na_2_^13^CO_3_.

**Fig. 2.**
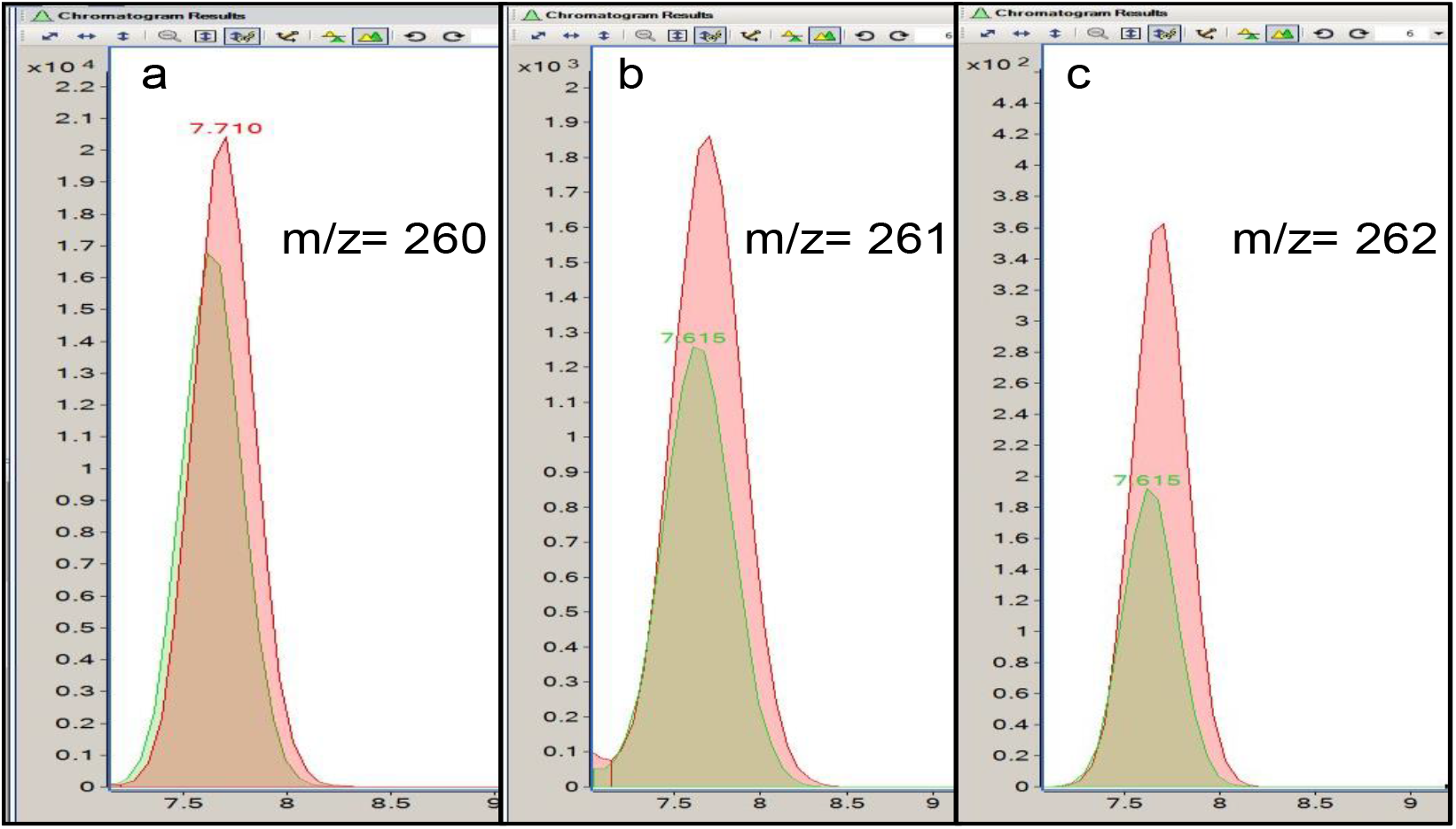
**(a-c):** The representative mass distribution of different mass isotopomers of Ala extracted from ^13^C-labelled (red) and unlabelled (green) leaf samples at 2 min after start of ^13^CO_2_. The greater peak area for isotopomers with *m/z* values of 261 (M+1) and 262 (M+2) compared to *m/z* of 260 (M) indicates higher ^13^C enrichment of Ala in ^13^C-labelled as compared to the unlabelled leaf sample.

To achieve ~ 400 ppm of ^13^CO_2_, 8 mL of 50 mM NaH^13^CO_3_ was placed in the Petri dish inside the container. The container was then placed on the stirrer plate to start rotating the magnetic bead. At the same time, IRGA was calibrated to zero for CO_2_ and H_2_O using a CO_2_ scrubber and desiccant, respectively. Other parameters of the IRGA were set as, photosynthetically active radiation (PAR), 1000 μmol m^−2^ s^−1^; flow, 500 μmol m^−2^ s^−1^; leaf and chamber temperature, 25°C. The CO_2_ scrubber was set to full bypass, while the desiccant was adjusted to maintain 55% relative humidity (RH) inside the leaf chamber. A fully expanded tobacco leaf was placed in the leaf chamber of the IRGA, and reference and sample values for CO_2_ and H_2_O were matched. After 5 minutes of stabilization, 1 mL of 50 mM HCl was added to NaH^13^CO_3_ to release ^13^CO_2_. At this point, a 5 seconds wait period was given to replace the existing ^12^CO_2_ with ^13^CO_2_ as detailed in previous studies (Xu et al., 2021, Evans et al., 2022). After the defined time intervals (30 seconds and 2 minutes), the leaf area outside the chamber was trimmed and immediately frozen in a container of liquid nitrogen placed below the leaf chamber of IRGA. The time lapse between the removal of trimmed leaf from the chamber to the freezing in liquid N_2_ should not be greater than 3 seconds. It is important to note that longer duration of ^13^CO_2_ feeding (>1 min) create vacuum in the external container which may result in error in the IRGA. To address this issue, the injection syringe can be removed while keeping the needle inserted in the top cover. The rear end of the needle should then be gently pressed with the thumb until the desired feeding time is reached. The feeding experiment was carried out with three independent biological replicates and the unlabelled leaves served as experimental control.

### Extraction and Derivatisation of amino acids

For extraction of free amino acids, leaf samples were crushed into fine powder in liquid nitrogen and suspended in 500 µl of 10 mM HCl. The clear supernatant obtained after centrifugation was collected as an amino acid extract. Amino acid derivatization was performed essentially as described by the manufacturer’s instructions (Waters, USA). Briefly, 10 µl of the extract/standard solution was mixed with 20 µl of aminoquinolyl-N-hydroxysuccinimidyl carbamate (AQC) and 70 µl of borate buffer followed by incubation at 55° C for 10 min. The mixture was filtered through a 0.25 µm filter before analysis.

### LC-MS Analysis

Derivatized amino acids were separated on an Agilent Infinity II UHPLC (Agilent Technologies, USA) and detected with a diode array detector (DAD) with separation column, ACCQ-TAG^TM^ ULTRA C18 (2.1 mm X 100 mm, 1.7µm). The water (A) and acetonitrile (B) each with 0.1% formic acid were used as a mobile phase in the maintained gradient system. A gradient elution method by varying the percentage of mobile phases A and B as 0-0.54 min, 99.90% A; 0.54-5.74 min, 90.0% A; 5.74-7.74 min, 78.80% A; 7.74-8.04 min, 40.40% A; 8.04-8.64 min, 40.40% A, 8.64-8.73 min, 99.90% A, 8.73-9.50 min, 99.90% A, 9.50-10.20 min, 99.90% A, and 10.20-11.00 min, 99.90% A, resulted in the separation of amino acids. Other UHPLC parameters were set as column temperature: 55 °C; injection volume: 2 µl; flow rate: 0.700 ml/min; and absorption wavelength of 260 nm.

The mass determination was performed using a high-resolution 6560 ion mobility quadrupole time-of-flight (IM-QTOF) LC-MS system (Agilent Technologies, USA) equipped with electrospray ionization (ESI) in positive mode. The drying gas flow rate was adjusted to 10.0 L/min, maintaining a constant gas temperature of 340 °C and a gas pressure of 35 psi. Additionally, the capillary voltage was set to 3500 V, accompanied by a fragmentor voltage of 390 V. The mass spectrometer scanned within the range of 100–1700 m/z at a rate of 1.50 spectra/s.

### ^13^C Enrichment analyses

The ^13^C enrichment in each amino acid was calculated as the percentage of ^13^C pool relative to the total carbon (both labelled and unlabelled) in the amino acid. Using mass spectrometry (MS) data, the enrichment was determined by calculating the percentage of the isotopomer peak areas containing ^13^C atom (M+1, M+2, M+3, etc.) relative to the total peak areas, including both labelled and unlabelled carbon (M+0, M+1, M+2, M+3, etc.). Additionally, the percentage change in the relative ^13^C enrichment between labelled and unlabelled control plants was calculated for each time interval (30 seconds and 2 minutes) of ^13^CO_2_ feeding.

## Results and Discussion

To validate the functionality of the ^13^CO_2_ delivery system, amino acid fraction obtained from ^13^C-labelled and unlabelled control plants were analysed using LC-MS. A total of eight amino acids (Ser, Gly, Asp, Glu, Thr, Ala, Pro, Cys) were successfully separated using LC-MS and subsequently analysed for ^13^C enrichment. It is important to note that for calculation, the mass of the derivatizing agent, AQC (171 Da), was added to the mass to charge ratio (*m/z*) of each isotopomer of the corresponding amino acid. For example, in the case of alanine (89 Da), the isotopomer containing only ^12^C will have a *m/z* of 260 (89 + 171 = 260) due to the addition of the derivatizing agent (AQC, 171 Da). The isotopomer with one ^13^C atom will have a *m/z* of 261 (90 + 171 = 261), while the isotopomer with two ^13^C atoms will have a *m/z* of 262 (91+ 171 = 262), and so on. Representative mass distribution of Ala extracted from ^13^C-labelled and unlabelled leaf samples at 2 min after the start of ^13^CO_2_ feeding is shown in the Fig. 2.

The data showed that after 30 seconds, only two amino acids, serine and alanine, exhibited significant increase in ^13^C enrichment, with percentages of 8.9% and 8.3%, respectively as comparted to the unlabelled control. On the contrary after 2 minutes, there is significant increase in the ^13^C enrichment in Ser, Gly, Asp and Ala by 24.1%, 13.8%, 13.5% and 25.9%, respectively as compared to the unlabelled control. No significant increase in the ^13^C enrichment was observed for amino acids, Glu, Thr, Pro, and Cys in ^13^C labelled and unlabelled plant samples (Fig. 3).

**Fig. 3:**
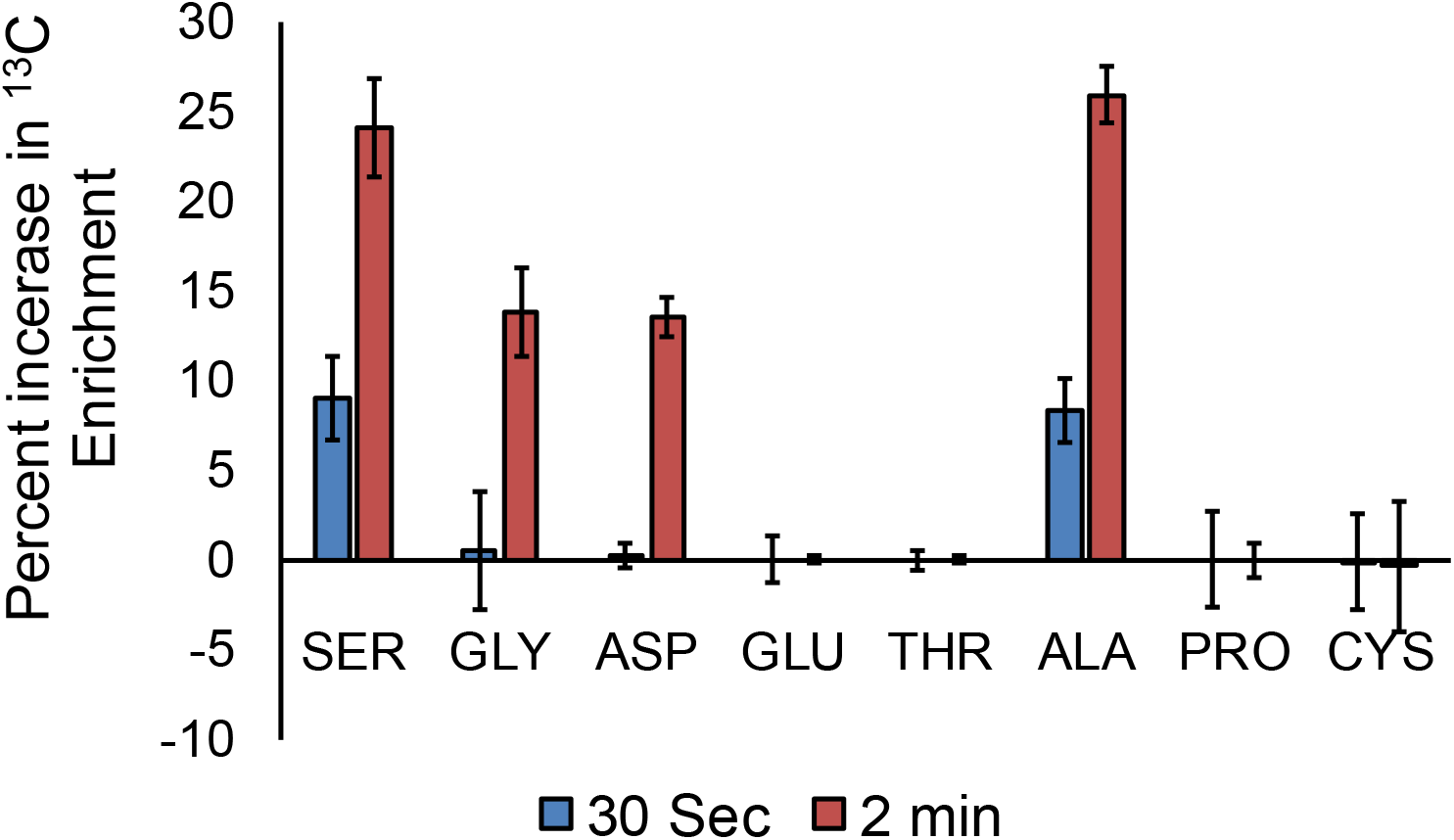
The percent increase in ^13^C enrichment in different amino acids after 30 sec and 2min of ^13^CO_2_ feeding as compared to the unlabelled control plants.

These results are consistent with a previous study, which demonstrated that Ser and Ala incorporated the ^13^C label in a short time interval after ^13^CO_2_ feeding, while Gly and Asp were labelled later in ^13^CO_2_-exposed *Arabidopsis* rosettes (Szecowka et al., 2013). Similarly, a study on the kinetics of photosynthetic ^13^C labelling in maize found that the percentage enrichment of ^13^C was higher in Ser, Gly, Asp, and Ala over the course of the labelling time scale (Medeiros et al., 2022), further validating our results.

It is worth noting that previous studies have utilized IRGAs as part of a ^13^CO_2_ delivery system. However, in these setups, the primary function of the IRGA is to monitor the concentration of ^12^CO_2_ inside the photosynthesis chamber. For instance, Evans et al. (2002) designed an improved dynamic flow cuvette for ^13^CO_2_ labelling in *Arabidopsis* plants, where the IRGA was used to track the replacement rate of ^12^CO_2_ with ^13^CO_2_ (Evans et al., 2002). In other studies, where the IRGA leaf chamber was employed for ^13^CO_2_ feeding, additional equipment, such as modified leaf chamber, an external CO_2_ cylinder or mass flow controllers, was required (Wright et al., 2014; Xu et al., 2021). The primary goal of the present study is to develop a simple ^13^CO_2_ delivery system using a conventional IRGA, aimed at making photosynthetic tracer studies more accessible. This system is portable, providing the added benefit of enabling ^13^CO_2_ feeding in plants growing in their natural habitat. Researchers often seek to conduct photosynthetic flux analysis under various environmental conditions, such as high altitudes, arid regions, and saline environments. The proposed system is particularly valuable in these studies, as transplanting plants from their natural habitat to laboratory conditions can lead to physiological and metabolic disturbances. Additionally, there are studies focused on understanding the flow of photosynthetic carbon across different plant types, including C3, C4, or C3-C4 intermediates, as well as in plants modified for enhanced photosynthesis through transgenic or genome editing techniques. In our view, by carefully implementing the ^13^CO_2_ delivery method and by using a larger number of replicates, the system described in the present work would help to draw a definitive conclusion on modulation of C flux with greater confidence.

## Acknowledgment

The Council of Scientific and Industrial Research (CSIR), India, financially supported this study through institutional grants (MLP-201).

## Data availability statement

The data that support the findings of this study are available from the corresponding author upon reasonable request.

